# Interplay between Rac1/RhoA and actin waves in giant epithelial cells : experiment and theory

**DOI:** 10.1101/2025.05.02.651882

**Authors:** Rémi Berthoz, He Li, Marie André, Michèle Lieb, Lucien Hinderling, Benjamin Grädel, Jakobus Van Unen, Olivier Pertz, Karsten Kruse, Daniel Riveline

**Affiliations:** Institut de Génétique et de Biologie Moléculaire et Cellulaire, Illkirch, France; Université de Strasbourg, Illkirch, France; Centre National de la Recherche Scientifique, UMR7104, Illkirch, France; Institut National de la Santé et de la Recherche Médicale, U964, Illkirch, France; Institute of Cell Biology, University of Bern, Baltzerstrasse 4, 3012 Bern, Switzerland; Department of Biochemistry, University of Geneva, CH-1211 Geneva, Switzerland; Department of Theoretical Physics, University of Geneva, CH-1211 Geneva, Switzerland

## Abstract

The acto-myosin cytoskeleton is a key driver of cellular shape changes *in vivo* and *in vitro*. Acto-myosin organization results from actin assembly and interactions between actin and myosin, which are both regulated by small Rho GTPases like Rac1 and RhoA. To uncover principles governing cytoskeletal organization, we analyzed actin patterns using live microscopy and theory. In giant Madin-Darby Canine Kidney (MDCK) epithelial cells and REF52 fibroblasts, we observed acto-myosin stress fibres and propagating waves. Stress fibres were stationary and correlated with homogeneous distributions of Rac and RhoA activity. Waves propagated at ≈ 1*μ*m/min and were associated with density variations of actin, Rac and active RhoA. Some waves transported cellular components or generated protrusions at the cell edge. Essential features of wave propagation are captured by a polar reaction-diffusion system for actin and Rac. Notably, two colliding waves annihilate each other. In cells, myosin activity was not required for the emergence of waves, but tended to suppress them. Consistently, local activation of RhoA slowed down or stopped and broke waves. These results highlight how the coupling between acto-myosin and Rho GTPase generates a variety of cytoskeletal structures and dynamics.

SIGNIFICANCE
The cytoskeleton shapes cells *in vitro* and *in vivo*. Its molecular mechanisms have been extensively documented, but the mesoscopic structures resulting from the interplay between the activator and the motor activity have been poorly characterized. Here we show that two structures are conserved in two systems, stress fibres with RhoA activity and diffusion actin waves regulated by Rac1. The annihilation of waves and the alteration of the wave dynamics are reproduced by a minimal model based on a polar reaction-diffusion model. This approach could pave the way to a generic description of regulated active gels in cells with potential biological functions, such as stress generation for fibres and probing of space and homogenization of the cortex by actin waves.

## INTRODUCTION

The actin cytoskeleton plays a crucial role in the determination of cell shape and drives essential cellular processes (1). To this end, the actin cortex forms specific structures such as stress fibres, lamellipodia and filopodia (2). Similar structures can be reconstituted *in vitro* (3, 4) and many of them rely on continuous actin turnover (5). It is thus probably not surprising that these cortical structures can be dynamic as illustrated by actin polymerization waves (6–9). Such waves have been observed in various cell types and play an important role in cell motility (6–8, 10, 11).

Cortical actin structures are often characterized based on their molecular composition (12). However, they arise from collective effects, that is, from interactions between many filaments, molecular motors, and cross-linkers. Also, they extend far beyond molecular scales. Physical descriptions have proven useful to analyze the mechanisms underlying the formation of cortical actin structures, for example, membrane ruffles (13, 14), stereocilia (15), and focal adhesions (16). These studies emphasize mechanical aspects and are carried out in the framework of active fluids, where the cytoskeleton is considered as a continuum and the action of molecular motors is captured by a so-called active mechanical stress (17).

In cells and in tissues, however, the cytoskeleton does not operate in isolation, but is coupled to cellular signalling. Notably, small Rho GTPases, such as RhoA, Rac1, and Cdc42, can orchestrate the dynamics of actin and myosin through their interaction with regulatory guanine nucleotide exchange factors (GEFs) and GTPase-activating proteins (GAPs) that activate and inactivate small GTPases (18). This regulatory framework is essential for the self-organization of the actin cytoskeleton into cellular structures (19, 20). Our understanding of how these regulatory mechanisms couple to the mechanical properties of the cytoskeleton remains limited, as such couplings are often overlooked in physical studies of active fluid systems (17). Notable exceptions in the theoretical treatment of regulated active gels are studies in the context of *Dictyostelium discoideum* chemotaxis, which focussed on the role of cytoskeletal regulation by membraneous phosphatidylinositol (3,4,5)-trisphosphate (PIP3), PI3-kinase, and IP3-degrading phosphatase PTEN (21, 22). These elements are downstream of the Ras family of small GTPases (23–26). Importantly, studies of the PIP3/PI3K/PTEN pathway revealed excitable dynamics of the actin cortex (27).

Theoretical investigations of active fluids coupled to regulatory biochemical networks have uncovered the potential to generate a variety of states that are absent in autonomous active fluids (28–32). In particular, such feedback provides a way to generate excitable dynamics (9, 33, 34) that is an alternative to the intrinsic excitability of signalling pathways. As we start to understand better the feedback from the cytoskeleton to the regulators (9, 35), it is possible to go beyond studying generic coupling terms and consider specific systems. At the same time, such a comparison requires experimental systems in which cytoskeletal structures can be studied, principally through imaging and mechanical perturbation. This requires systems of a large size. Examples are *Xenopus* oocytes (9), starfish oocytes (36), and giant *D. discoideum* cells (27).

To investigate the dependence of acto-myosin self-organization on regulation by small Rho GTPases, we developed an assay to quantitatively characterize cortical actin patterns and their dynamics. Specifically, we imaged the dynamics of actin in giant Madin-Darby Canine Kidney (MDCK) epithelial cells of typically 100μm in diameter (37). This approach provided a large surface area for the emergence and evolution of cytoskeletal structures away from cell-cell junctions. Within the giant cells, we observed fibres, localized states, and waves. Actin fibres existed in a homogenous background of active RhoA. The waves depended on actin assembly and activity of Rac1, which also formed a travelling wave. Remarkably, these polymerization waves drove transport processes. We also report similar structures in REF52 fibroblast cells which points towards the generic nature of these phenomena. We quantitatively characterized the waves and found that they are well described by a reaction-diffusion system. The results of our study show that salient cortical actin structures result from mutual feedback between the spatio-temporal dynamics of the cytoskeleton and small GTPases.

## MATERIALS AND METHODS

### Plasmids and cell lines

To manipulate Rac activity in single living cells, we re-engineered an iLID-based optogenetic actuator based on a DH/PH domain of the GEF Tiam (38). We used a high-affinity nanoSspB domain for efficient light-dependent recruitment of the Tiam GEF domain to the iLID anchor. To better focus optogenetic activation, we also fused the iLID anchor to a stargazin membrane anchor that slows its diffusion (39). The stargazin-iLID anchor and SspB-LARG domains were separated by a self-cleaving P2A peptide, allowing equimolar expression of both units from a single operon. We refer to this construct as optoTiam. We developed a stable MDCK II cell line transfected with LifeAct::iRFP703 as an actin reporter. In addition, we introduced other reporters depending on the experimental requirements. We prepared a stable cell line expressing MRLC::GFP (myosin reporter) and 2xrGBD::dTomato (RhoA activity biosensor). To visualize Rac1 dynamics, we used transient transfection with Rac::GFP. For optogenetic experiments, we generated stable cell lines expressing iLID-LARG::mVenus (RhoA-GEF optogenetics) and 2xrGBD::dTomato, as well as iLID-TIAM::mVenus (Rac-GEF optogenetics). We also used REF52 transfected cells and their cytoplasts (40).

### Cell culture

Cells were maintained at 37 °C in a 5 % CO2 incubator with 85 % humidity. They were cultured in the following culture medium: Minimum Essential Medium with Glutamax-I with 5 % of Fetal Calf Serum, 1 % Non-Essential Amino Acids, 1 % Sodium Pyruvate and 1 % Penicilin-Streptomycin. Every 2 to 3 days, cells were resuspended and diluted in order to maintain the culture below 80 % confluency. Resuspension was performed by detaching cells from the Petri dish using Trypsin/EDTA after a wash with PBS. The suspension was centrifuged at 300 g for 2-3 min. Cells were resuspended in culture medium, their concentration measured, and the proper dilution was used in order to seed cells at a density of 400 cells/cm^2^ to 4000 cells/cm^2^ depending on experiments.

### Preparation of giant cells

Giant MDCK cell samples were produced by seeding a freshly resuspended culture of MDCK cells, with 0.5 μM of cytochalasin-D (Sigma-Aldrich, C8273, St. Louis, MO) in the culture medium (37, 41). In preparation for microscopy, they were seeded directly on a microscope glass. To minimise cell-cell interactions they were seeded at a density of 1500 cell/*cm*^2^. Cells were maintained in the incubator; the culture medium was replaced every 2 to 3 days. Samples were used between 3 and 7 days of incubation with cytochalasin-D with a typical diameter that increased with the incubation time (Supp. Fig. 1). Cytochalasin-D was washed-out before between 1 and 5 hours before acquisition.

### Immunofluorescence microscopy and dyes

We checked that our transfections did not affect the formation of patterns by labelling actin and myosin using immuno-fluorescence microscopy (42). Briefly, we fixed cells with paraformaldehyde and we permeabilised cells with Triton before incubation with primary and secondary antibodies. We used the following labels and antibodies: DAPI (Sigma-Aldrich, MBD0015, St. Louis, MO), Phalloidin::Alexa405 (Invitrogen, A30104, Waltham, MA), Phalloidin::Alexa488 (Invitrogen, A12379, Waltham, MA), Rabbit-MLC2 (Cell Signaling, 3672, Danvers, MA), Goat anti-rabbit::Alexa647 (Invitrogen, A-21245, Waltham, MA). We checked that cytoskeletal structures were similar to the dynamical structures found in our transfected cells (Supp. Fig. 2). For Fig. 2, we labelled the cell membrane with the MemGlow::488 membrane dye (Cytoskeleton, MG01-02, Denver, CO) at a concentration of 40 nM.

**Figure 1:**
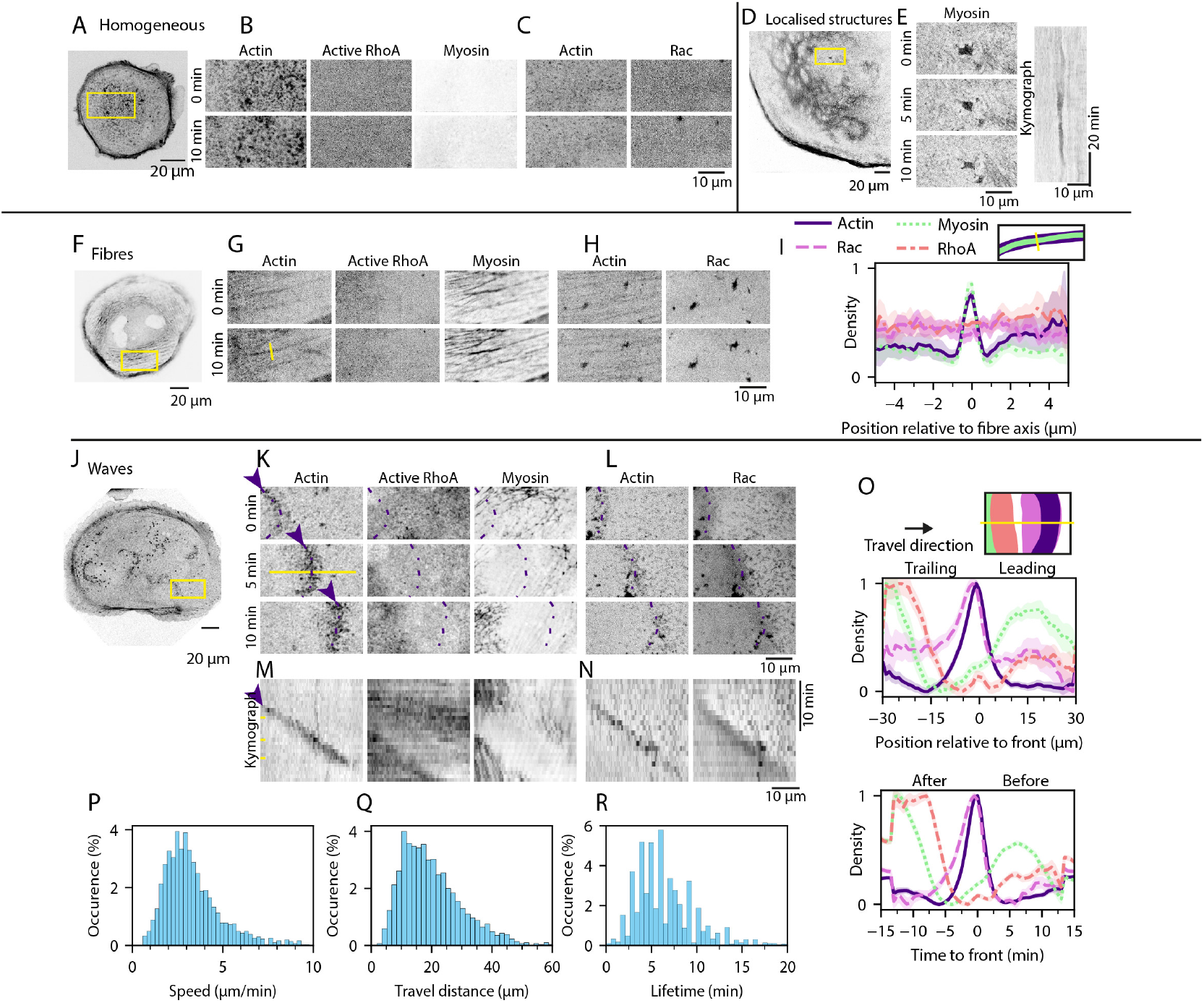
The cell cortex displays four kinds of organised structures in giant MDCK cells. A-C: Homogeneous cortex (Movie 1). A: LifeAct-RFP. B: Snapshots of the region of interest (ROI) indicated by yellow rectangle in A showing LifeAct-RFP (Actin), GBD-dTomato (active RhoA), and MRLC-GFP (Myosin). Signals are homogeneous in space and do not evolve into patterns within 10 minutes. C: Snapshots of an ROI in a different giant MDCK cell, showing LifeAct-RFP (Actin) and Rac-GFP (Rac). D, E: Localised structure (Movie 2). D: MRLC-GFP. E: Time series of the ROI (yellow rectangle) in D (left) and kymograph (right), showing the long-term stability of the localised structure. F-I: Fibres (Movie 3). F: MRLC-GFP. G: Snapshots of ROI (yellow rectangle) in F, showing LifeAct-RFP (Actin), GBD-dTomato (active RhoA), and MRLC-GFP (Myosin). Actin and myosin signals show fibres persisting over 10 minutes, active RhoA shows no structure. H: Snapshots of an ROI in a different giant MDCK cell, showing LifeAct-RFP (Actin) and Rac-GFP (Rac). I: Quantification of actin, myosin, Rac and active RhoA densities along the yellow line in G. Data averaged from *N* = 3 biological repeats, *n* = 9 cells, *w* = 174 fibres. J-O: Waves (Movies 4 and 5). J: LifeAct-RFP. K: Snapshots of the region of interest (ROI) indicated by yellow rectangle in J showing LifeAct-RFP (Actin), GBD-dTomato (active RhoA), and MRLC-GFP (Myosin). A front of actin (violet arrow heads) travels from left to right. Violet dashed lines indicate position of the actin front. There are no myosin or active RhoA waves. L: Snapshots of an ROI in a different giant MDCK cell, showing LifeAct-RFP (Actin) and Rac-GFP (Rac). Rac shows a wave moving with actin. M, N: Kymographs from the time series in K, L. O: Quantification of actin, myosin, Rac and active RhoA densities along the yellow line in K. Myosin and active RhoA are present in front of and behind the actin wave, but absent in region of high actin and Rac density. Actin density increases first, Rac density is high in the wake of the wave. The 4 densities as a function of time at a fixed point in space. The passing wave first exhibits a decrease of myosin density, followed by an increase of the actin and then the Rac density. After the wave front has passed, RhoA activates and myosin density increases. Data for actin, myosin, RhoA activity averaged from *N* = 2 biological repeats, *n* = 14 cells, *w* = 239 waves, and for Rac averaged from *N* = 2, *n* = 10, *w* = 58. P-R: Quantification of wave properties: wave speed (P), travel distance (Q) and lifetime (R). Data cumulated from *N* = 10, *n* = 98, and *w* = 1528.

**Figure 2:**
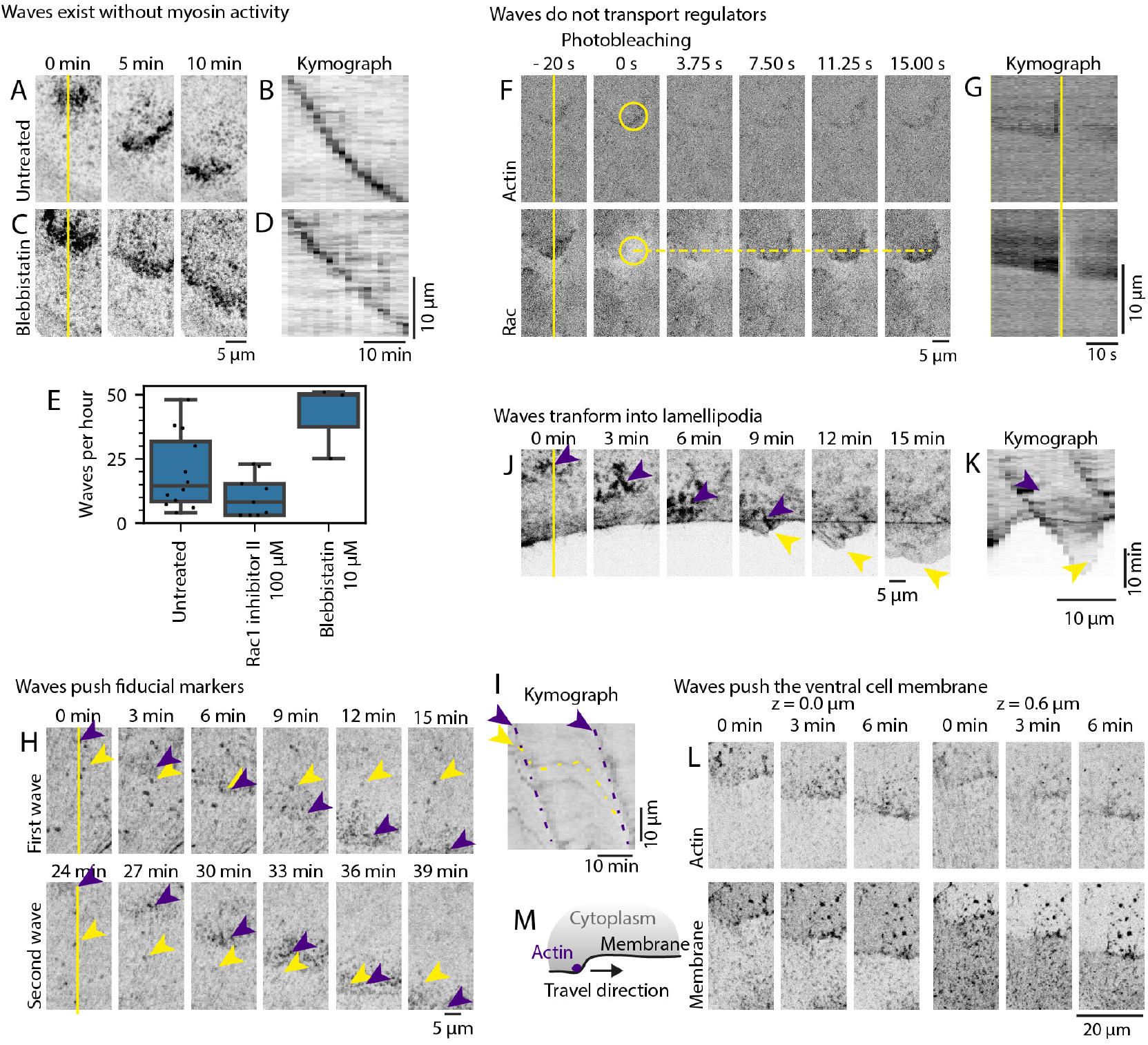
Waves are actin- and Rac-dependent and have mechanical impact in giant MDCK cells. A: Time series of LifeAct-RFP (Movie 10). B: Kymograph along the central vertical line of A, indicated by the yellow line. Unless otherwise specified, all the following kymographs are sampled along the center of the images. C: Time series of LifeAct-RFP in the same cell and at the same location as in A after 2 hours and addition of 10 *μ*M blebbistatin (Movie 10). D: Kymograph along yellow line in C. E: Number of waves per hour in untreated cells (*N* = 10 biological repeats, *n* = 98 cells, *w* = 1528 waves), in the presence of 100 *μ*M Rac inhibitor II (*N* = 3, *n* = 33, *w* = 320, = 2.7 10^−2^), and 10 *μ*M blebbistatin (*N* = 3, *n* = 33, *w* = 747, *p* = 2.5 · 10^−1^). P-values from Wilcoxon test between drug and untreated condition. F: Fluorescence recovery after Rac-GFP photobleaching in yellow circle at time 0 s (Movie 13). LifeAct-RFP (Actin) and Rac-GFP (Rac). Rac fluorescence recovers within seconds. Yellow dashed line indicates the position of the wavefront at time −20 s. G: Kymographs from time series in F. Yellow line indicates time 0 s. H: Time series of LifeAct-RFP for two subsequent waves. Violet arrow heads indicate the wave fronts, yellow arrow heads a fiducial marker (Movie 14). The marker is moved by the passing wave and (partially) relaxes after the passage. I: Kymograph from the time series in H. Violet dashed lines: trajectory of wave fronts, yellow dashed line: trajectory of the fiducial marker. J: Time series of LifeAct-RFP. Violet arrow heads indicate a wave front. As the front reaches the cell edge, a lamellipodium is generated (yellow arrow head, Movie 16). K: Kymograph from the time series in J. Violet arrow head: trajectory of the wave front, yellow arrow head: edge lamellipodium. The extension speed of the lamellipodium matches the wave propagation speed. L: Time series of LifeAct-RFP and membrane in 2 different *z*-sections. M: Schematic representation in the *xz*-plane of the data in L.

### Microscopy and live cell imaging

Samples were mounted in an incubation chamber at 37°C with 5% CO2 and 85% humidity and cells were maintained in culture medium. Samples were imaged with either a spinning disk or a scanning confocal fluorescence microscope. The spinning disk was a Leica (Wetzlar, Germany) DMi8 microscope equipped with a Yokogawa (Tokyo, Japan) CSU-W1 confocal system using a Leica (Wetzlar, Germany) Plan-Apochromat 63X/1.4 NA oil objective and a Hamamatsu (Bridgewater, NJ) camera. Live imaging was performed with a time interval of 30s or 1min. The microscopes were controlled with the MetaMorph software. The scanning confocal was a Leica (Wetzlar, Germany) SP8-UV microscope equipped with an Argon laser, a HeNe laser and a 561 nm laser diode, using a Leica (Wetzlar, Germany) Plan-Apochromat 40X/1.3 NA oil or 63X/1.4 NA oil objective. Live imaging was performed with a time interval of 30 s or 1 min, cells samples were mounted in a temperature-controlled chamber at 37 °C and culture medium was replaced with Leibovitz’s L-15 medium with 5 % Fetal Calf Serum and 1 % Penicilin-Streptomycin. The microscope was controlled with Leica’s LAS X software. Optogenetic experiments were performed on the scanning confocal microscope. The scanning module was used to illuminate the sample within one or more restricted regions of interest (ROI) of the field of view and within a restricted period of time (using Leica’s LAS X module Live Data Mode). The module was configured to illuminate the ROI every 10 s during the stimulation period, using the 458 nm laser at 10 % power and 600 ns dwell time. Fluorescence frames are acquired in-between stimulations, every 30 s or 1 min. FRAP experiments were performed on the spinning disk confocal microscope. For these experiments, the image time interval is set to 1.17. Photobleaching was performed with the same illumination sources as for acquisitions, but the power is set to 80 % and exposure set to 1 s.

### Drug treatments

In some experiments, cells were treated with specific inhibitors at the indicated concentrations. We used blebbistatin (Sigma, B0560) at concentrations of 10 μM or 20 μM and Rac1 Inhibitor II (InSolution, Z62954982) at a concentration of 100 μM. The inhibitor was added in the culture medium during live imaging at the indicated time points. In experiments with blebbistatin, the 488nm illumination channel is turned off to avoid phototoxicity.

### Image analysis and statistics

All analyses were performed with the Fiji (43) software for image visualisation and measurements, and custom Python scripts for measure processing and graphing. To measure protein density profiles in wave fronts and wave speed, we plotted kymographs using the custom-made ‘Live Kymographer’ Fiji plugin. To draw kymograph of waves, we used the actin fluorescence recordings as a readout for wave localisation. Kymographs were drawn parallel to the direction of wave propagation, in single straight lines, as long as the wave propagated in a single direction. Waves that changed direction were measured as multiple independent waves if all directions were significant in terms of traveled distance, or were measured only once otherwise. Waves that expanded radially were measured only once, not in multiple directions. The wave speed was directly computed with a Python script as the ratio of distance of travel to duration of travel. To extract protein density profiles in wave fronts from kymographs, we translated the reference frame of the kymograph from the microscope reference frame to the reference frame co-moving with the wave, using a Python script. In this reference frame, spatial protein density profiles were essentially constant over time. For each wave and each fluorescence channel, we averaged the spatial profiles over time with a Python script to get a one-dimensional (spatial) profile of the wave. For analysis involving multiple waves (*i*.*e*. averaging of multiple waves, alignment of multiple fluorescence channels, …), wave profiles were aligned based on their actin profile which defines the wave front. The number of experimental repeats is indicated in figure captions, with N for biological repeats, n for number of cells, w for number of waves or fibres.

### Reaction-diffusion equations for the interplay between actin assembly and Rac1

We describe the interplay between actin assembly and Rac1 using a previously developed set of reaction-diffusion equations (33, 44). In this approach, the state of the system is captured by the actin density *c*, the local polarization vector **p**, and the densities *n*_*a*_ and *n*_*i*_ of active and inactive Rac1. The dynamic equations are given by

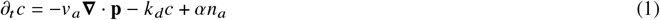

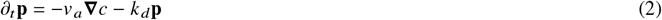

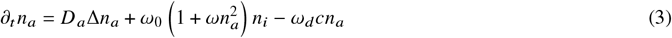

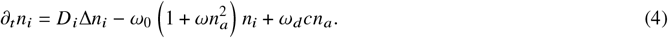

where *v*_*a*_ is the polymerisation velocity of actin, *ω* the spontaneous rate of Rac1 activation, *ω*_0_ governs the rate of auto-catalytic Rac1 activation, and *k*_*d*_ quantifies actin-induced Rac1 inactivation. Furthermore, *α* is the rate of actin assembly induced by active Rac1 and all species diffuse with respective diffusion constants *D, D*_*a*_, and *D*_*i*_. Note that the total number of Rac1 is conserved ∫_*A*_(*n*_*a*_ + *n*_*i*_)d*A* = *An*_total_ = const.

We solve the equations using a semi-spectral method and on a square grid of size 256 × 256 with periodic boundary conditions for space. To do this, we use the code published in (45), where time is given in units of 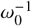 and length in units of 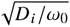. We use a time step = 10^−6^. The parameter values are *v*_*a*_ = 0.7, *ω*_*d*_ = 0.35, and for all other parameters as in Ref. (33). To describe Rac1 inhibition, we reduce the Rac1 activation rate by 25% in a circular region of radius *R* = 16.

As initial condition, we chose in all cases *c* = 0 and **p** = 0. In Fig. 3A, we initially set *n*_*a*_ = 170 and *n*_*i*_ = 1 in a circular region of radius *R* = 16, whereas *n*_*a*_ = 1, *n*_*i*_ = 170 elsewhere. To simulate wave annihilation in Fig. 3C, two wave sources are separated by 64 in our computation.

**Figure 3:**
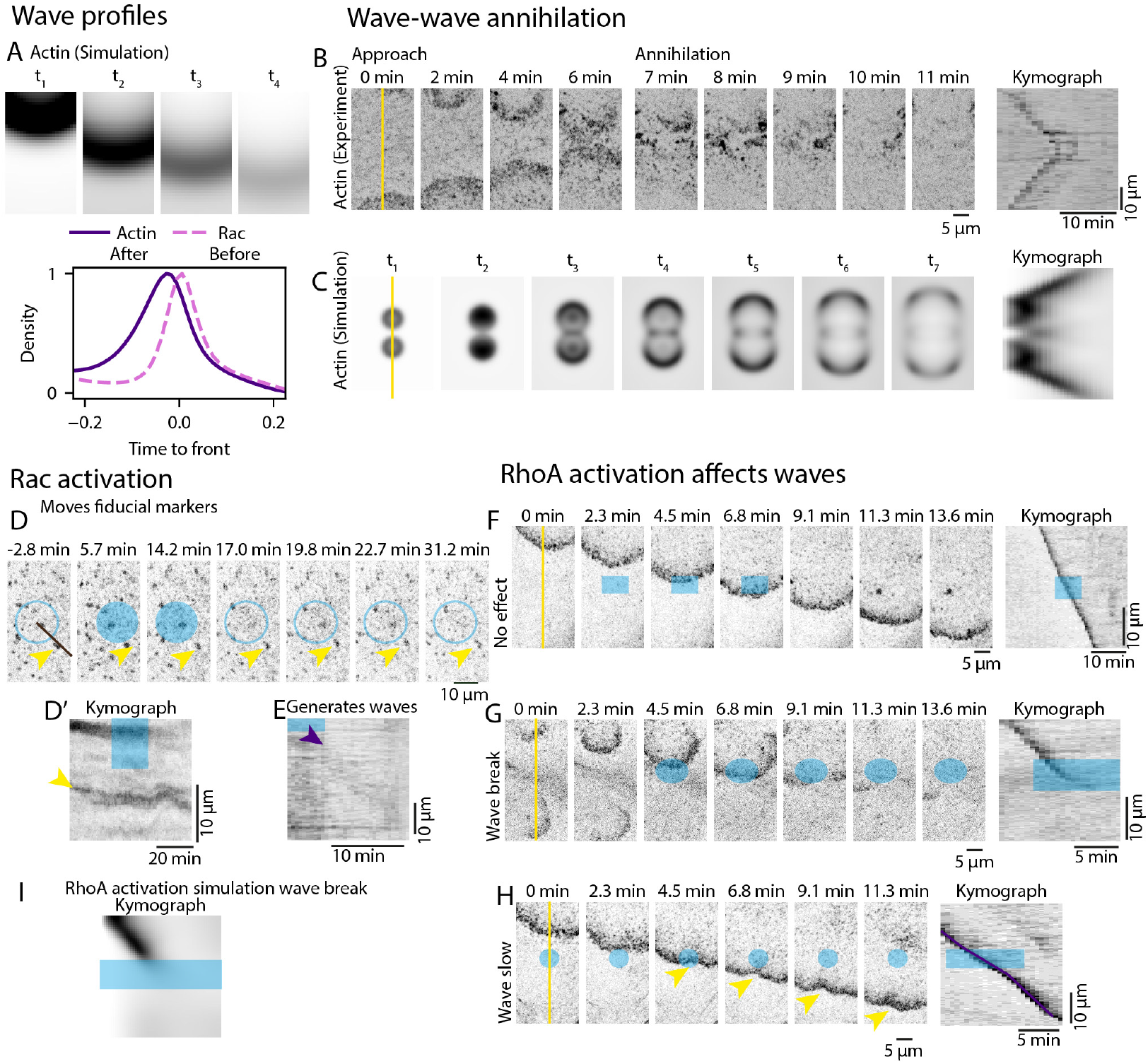
A polar reaction-diffusion system recapitulates features of wave propagation A, top: Snapshots of actin and Rac densities for a wave from a 2d numerical solution to the polar reaction-diffusion system (1)-(4) (Movie 17). Bottom: Actin and Rac profiles along the direction of wave motion. B, left: Snapshots of LifeAct-RFP a giant MDCK cell showing two waves approaching and colliding (Movie 20). Right: kymograph along the yellow line in snapshot at *t* = 0 s. C, left: Snapshots of actin density for two colliding waves from a 2d numerical solution to Eqs. (1)-(4) (Movie 18). Right: kymograph along the yellow line in snapshot at *t* = *t*_1_. D: Optogenetic activation of Rac. Snapshots of Rac-GFP density in a giant MDCK cell. Between *t* = 0 min and *t* = 15 min, Rac1 is activated in the blue region. A fiducial marker (yellow arrow head) moves during stimulation and relaxes afterwards (Movie 21). D’:Kymograph of Rac-GFP along the black line in D, blue region indicates Rac1 activation. Yellow arrow head indicates the fiducial marker. E: Kymograph of Rac during and after optogenetic activation of Rac1 in blue region (Movie 22). Upon release of stimulation, a wave is generated (violet arrow head). F-H, left: Snapshots of LifeAct-RFP in cells with optogenetic activation of RhoA (blue regions, Movies 23-25). Rac activation can have no effect, slow down or stop actine waves. Right: Corresponding kymographs along the yellow lines. I: Kymograph from a numerical solution to Eqs. (1)-(4), where RhoA activation is captured by Rac inhibition (Movie 19).

## RESULTS

### Characterisation of acto-myosin structures, regulation and dynamics

In this section, we provide an overview of the acto-myosin network structures formed in giant MDCK cells. We relate them to the distributions small GTPases, specifically active RhoA and Rac1.

#### Acto-myosin patterns in giant MDCK cells

To identify the major acto-myosin patterns, we prepared giant epithelial MDCK cells with a diameter of typically 100 μm. By using giant cells, we reduced the influence of the cell edge on the acto-myosin patterns that formed. They could thus be attributed to intrinsic properties of the network. Furthermore, structures with an intrinsic length scale of up to 10 μm could develop in these cells. To visualize filamentous actin and non-muscle Myosin II (myosin for short), MDCK cells were transfected with LifeAct-RFP and MRLC-GFP (Methods).

In cells at an early stage of spreading after cytoD washout, acto-myosin did not feature any prominent structure on the μm-scale (Fig. 1A-C and Movie 1). We call this state the homogenous state. Actin exhibited some granularity on sub-μm scales (Fig. 1A,B). With time, actin structures fluctuated, which indicated that filamentous actin turned over on a minute time-scale. In contrast, the myosin density showed less temporal and spatial fluctuations.

In some cells, myosin formed clusters with a size of roughly 1 μm and a life time of tens of minutes (Fig. 1D,E and Movie 2). These clusters were isolated and surrounded by regions with a homogenous myosin density. These clusters could be instances of so-called localized states that have been reported as solutions of physical descriptions of acto-myosin networks coupled to biochemical regulators (31).

Stable linear structures were a frequent acto-myosin pattern in our giant MDCK cells (Fig. 1F-H, Movie 3, Supp. Fig. 3). They spanned tens of micrometers and persisted for more than an hour. Due to the simultaneous high density of filamentous actin and myosin, we identified these structures as being stress fibres. Stress fibres were distributed non uniformly in space. In the region outlined in yellow in Fig. 1F, stress fibres were aligned and oriented along the cell boundary. Directly adjacent to stress fibres, the actin and myosin densities were below their background values (Fig. 1I).

The final pattern we recurrently observed, were travelling actin waves (Fig. 1J-L, Movies 4, 5, Supp. Fig. 3). Kymographs show that the wave propagation velocity was essentially constant (Fig. 1M, N). The waves propagated at an average velocity of (3.6 ± 1.4) μm min^−1^ (Fig. 1P), over a typical travel distance of 20 μm (Fig. 1Q), and had a typical life time of (5.8 ± 2.8) min (Fig. 1R). The lateral extension was typically 20 μm. Initially, waves commonly expanded radially before propagating with a fixed lateral length (Movies 4, 5).

These patterns were not mutually exclusive as some cells presented simultaneously, for example, stress fibres and homogenous regions. In addition, waves could propagate in regions containing stress fibres. All these patterns were essentially two-dimensional as they were present in flat regions of the cortical layer (Supp. Fig. 4).

#### Waves and fibres are also present in fibroblasts

Next, we tested whether similar structures were also present in fibroblasts. Similarly to giant MDCK cells, we also observed fibrous actin patterns in REF52 cells and in REF52 cytoplasts, where the nucleus had been removed (Supp. Fig. 5). Local contraction could be induced reversibly both in REF52 cells and their cytoplasts (Movies 6, 7). In addition, we observed spontaneous actin waves on the ventral surface of REF52 cells (Supp. Fig. 6 and Movie 8). We could even generate waves in REF52 cells by locally activating RhoA on the dorsal surface of the cells (Supp. Fig. 7, Movie 9).

#### Waves but not stress fibres are associated with patterns of RhoA or Rac1

We studied the regulation of actin assembly and acto-myosin contractility for generating the above patterns by imaging the distribution of active RhoA and of Rac1. Neither active RhoA nor Rac1 displayed heterogeneities across stress fibres (Fig. 1I), suggesting that these small GTPases are not involved in their maintenance.

This was different for actin waves (Fig. 1O). At a fixed time point, the high actin density region of a travelling wave co-localized with a depletion zone of active RhoA. The active RhoA density was higher behind the region of high actin density than in front. Similarly, myosin was depleted in the region of high actin density. Similar to active RhoA, the myosin density was higher behind the wave than in its front. Compared to the density of active RhoA, the myosin density was shifted towards the trailing end of the wave. In contrast to active RhoA, the density of Rac1 was increased in the region of high actin density with both maxima essentially coinciding.

### Waves are reaction-diffusion waves

In this section, we explore the mechanism underlying the propagation and emergence of actin waves.

#### Waves depend on actin polymerisation regulated by Rac1

The decrease of the myosin density in the region of high actin density in the waves points towards a gradient in active stress. With the myosin density being higher in front of the wave than in the region of high actin density, this could suggest that the actin network is pulled forward by the molecular motors. We tested the contribution of myosin contractility to the propagation of actin waves by applying the small molecule inhibitor of myosin blebbistatin. After treatment with blebbistatin, actin waves persisted (Fig. 2A-D). Furthermore, the number of waves emerging in a cell per time interval increased (Fig. 2E, Movie 10 and Movie 11), whereas their velocity decreased by 40 % (Supp. Fig. 8). In contrast, the stress fibres disappeared after incubation with blebbistatin (Movie 11).

The effect on actin waves of inhibiting Rac1 was different (Movie 12). In cells treated with Rac1 inhibitor II, the average number of waves in a given time interval decreased by 50 % (Fig. 2E) and their velocity by 20 % (Supp. Fig. 8). These findings suggest that actin waves are a consequence of Rac1-regulated actin assembly.

Since activation of small GTPases requires binding to the cytoplasmic membrane (18), the waves are localized to the cytoplasmic membrane. There are two general ways to produce protein waves on surfaces. Either the wave forming protein is displaced along the membrane while being bound to it. This is, for example, the case for anterior Par proteins during symmetry breaking at the one-cell stage of nematode embryos (46). Alternatively, the wave-forming proteins continuously shuttle between the membrane and the cytoplasm in such a way that they attach at the wave’s leading edge and detach at its trailing edge. An example of such waves are provided by the Min-proteins of *Escherichia coli*. The latter mechanism is reminiscent of waves in excitable media (33).

To evaluate the coupling between Rac1 and actin and determine whether Rac1 is transported, we performed FRAP experiments to locally bleach Rac1 (Fig. 2F, Movie 13). We observed a rapid recovery within seconds and concluded that the bleached regions did not move, suggesting that Rac1 is not transported along the membrane.

Next, we investigated whether mechanical function was associated with wave propagation (Fig. 2H-L). By tracking fiducial markers, we observed their motion, which correlated in both velocity and direction with wave propagation (Fig. 2H-I, Movies 14, 15). Additionally, we tracked membrane displacement, which showed an overshoot along the *z*-axis, corresponding with wave movement (Fig. 2L). Finally, we examined wave propagation at cell borders and observed their transformation into lamellipodia, protruding from the cells (Fig. 2J-K, Movie 16). Collectively, these results suggest that waves are dependent on actin and Rac1, with a mechanical impact on the environment.

### Actin waves are captured by a polar reaction-diffusion model

Based on these observations, we turned to theory to better understand wave behaviour. In a cell, the regulation of actin assembly involves a number of different proteins connecting Rac1 to actin promoting factors like formins or the Arp2/3 complex. Details of the coupling between the acto-myosin network and small GTPases regulating have been worked out in a few cases (35, 47). Theoretical descriptions of the cytoskeleton and regulators of its assembly and the active mechanical stress it generates have been introduced in Refs. (28–31, 48, 49). Similar to these works, here, we do not aim to describe all the intermediate steps. Instead, we content ourselves with an effective description, where all protein species involved in the regulation of actin assembly are effectively captured by the regulator (33). It accounts for the actin density and the alignment of the actin filaments as well as for a regulator of actin assembly. This regulator can be in an active and an inactive form and we refer to it with a slight abuse of language as Rac1. In brief, active Rac1 assembles new actin filaments, which in turn inactivate Rac1. Inactive Rac1 is activated spontaneously and cooperatively. Our description does not include Myosin-generated stress or flows as RhoA does not seem to be involved in wave generation. Given the fast recovery of Rac1 in the FRAP experiment (Fig. 2F), we also neglect a potential advection of Rac1 with acto-myosin flows. The system is defined in detail in the Methods.

We solved the dynamic equations numerically, explored the resulting phase diagram, and compared the solutions with experimental observations. We identified a set of parameters that closely matched the experimental data and used it to test variations in initial conditions and spatial perturbations (Supp. Mat.). An example of a travelling pulse is shown in Fig. 3A (Movies 17-19). The profiles of active Rac1 and actin peak sharply along the propagation direction, where active Rac1 precedes actin. Pulses that are initiated at a point, spread radially and remain circular symmetric. The peak densities decrease as the pulse spreads. These profiles are qualitatively similar to those observed experimentally (Fig. 1M). Note that in our experiments, Rac1 persists somewhat longer than actin, which is not the case in our theory. We ascribe this difference to intermediate steps in the negative feedback of actin on Rac1, which are not accounted for in our theory and which would delay Rac1 inactivation.

In our experiments, actin waves annihilated each other when colliding (Fig. 3B and Movie 20). This is consistent with previous observations of actin waves in other cell types (8, 27). It points to excitability of the actin cortex (27, 34, 50). Similarly, in our theory, two colliding wave fronts tend to annihilate (Fig. 3C) and the dynamic equations have been related to the FitzHugh-Nagumo system (33).

### Activation of Rac1 and RhoA affects wave propagation

Next, we examined how local activation or inhibition of Rac1 affected wave propagation (Fig. 3D-I). Optogenetic activation of Rac1 induced a local reversible expansion of the cortex as indicated by the displacement of fiducial markers (Fig. 3D, Movie 21). After stopping Rac1 activation, the cortex relaxed as shown by the motion of fiducial markers and a wave was generated (Fig. 3D’, E, Movie 22).

Even though inhibition of Myosin II did not affect wave propagation, optogenetic local activation of RhoA could have an effect on waves. In some cases, waves propagated unaffected through regions of activated RhoA (Fig. 3D, Movie 23). Also, we observed a slow down of wave propagation (Fig. 3F, Movie 24) and in the remaining cases waves broke as the part of the wave from traversing the activated area disappeared (Fig. 3E, Movie 25). We ascribe the effects of RhoA activation to a negative feedback of RhoA on Rac1 activity (35, 47). Indeed, we had observed a decrease in RhoA activity in regions of peak actin and Rac1 densities of travelling waves (Fig. 1M). When locally inactivating Rac1 in our theory, we observed breaking of a wave front (Fig. 3G).

## DISCUSSION

In this work, we have reported on cortical actin structures together with the distribution of regulating small GTPases. To reveal the internal dynamics of this system we generated giant cells. We found fibres, localized states, and waves, and investigated the dynamics of waves in detail. Our results show that a characterization of cortical actin structures requires a simultaneous analysis of small GTPases. As a consequence, physical descriptions of cortical structures, which have already provided important insights, also need to be amended to account for the coupling between active mechanics and biochemical regulation.

It is interesting to consider the potential functions of each structure in the light of the coupling between small GTPases and the actin cortex reported above. Fibres are known to generate local stress, to reinforce adhesion, and to contribute to cell motion (51). The homogenous and constant distribution of active RhoA in regions, where stress fibres are present, may play a crucial role in maintaining a constant stress that the cell applies to its environment. It also shows that the formation of stress fibres are inherent to the cortical actin dynamics and does not require signalling through small RhoGTPases. However, notably the formation of focal adhesions is still coupled to biochemical regulation, for example, through the Src family of tyrosine kinases.

In contrast to the fibres, the actin polymerization waves are accompanied by similar patterns in Rac1. This is reminiscent of actin-active RhoA waves reported in frog and starfish oocytes (9). Whereas actin polymerization waves might serve as guidance cues during cell migration (11), their function in oocytes is currently unknown. We speculate that these waves are involved in maintaining the integrity of the actin cortex and serve, in particular, to repair possible cortical ruptures. Dynamic regulator patterns can help to avoid amplification of assembly heterogeneities.

In tissues, actin waves are associated with the generation of mechanical stress (52). In this context, waves can be instrumental to direct tissue deformation during development, for example in *Drosophila melanogaster* (53). As these waves involve mechanical stress, the underlying mechanism is likely different from the reaction-diffusion waves we have considered above. To analyze the physics of mechanical waves, the above description needs to be generalized to also account for active stress generation by the cortical actin. Such generalized descriptions have revealed states like “fibre networks” (30) or localized states (31, 32) that need to be studied further experimentally. Similar approaches might also be useful to further investigate the emergence of stress fibres or the dynamics of focal adhesions as alluded to above. Beyond these, also the connection of membrane ruffles and retrograde flows to biochemical signalling can be studied in this context.

In our assay, we show that the regulators are not transported with the waves, Rac1 in waves spontaneously generated in giant cells, and RhoA in waves promoted by optogenetics in Ref52 cells. This result is in contrast with transport of small Rho GTPases reported *in vivo* in Drosophila (54), but is consistent with the absence of transport report in C. elegans in early stages of development (29). It will be interesting to test this transport in other situations in vitro and *in vivo* in order to evaluate its potential consequences. In particular localised states were predicted when regulators were transported and this could represent a paradigm for coupling between mechanics and regulation with new explanations for patterns formation of the cortex viewed as an excitable medium with mechanical actions.

## CONCLUSION

Our work reported structures regulated by signalling with an emphasis on waves and fibres. It will be interesting to test more GTPases such as Cdc42 for example and their interplay with their respective GEFs and GAPs on the mesoscopic features and their dynamics. Future developments in probes and in theory will allow to evaluate patterns dynamics and their implications during development. The level of integration of the regulated active gel allows to generate patterns with generic features prone to generic comparisons with experiments.

## AUTHOR CONTRIBUTIONS

DR and KK designed the research. RB performed experiments with support from MA, ML, LH, BG, JVU, OP. HL performed simulations. DR and KK wrote the article with inputs from all authors.

## ACKNOWLEDGEMENTS

We thank Luca Barberi (University of Geneva) and Damian Brunner (University of Zurich) for discussions and the Riveline team for feedback and suggestions. We thank E. Grandgirard and E. Guiot and the Imaging Platform of IGBMC. D.R. acknowledges the Interdisciplinary Thematic Institute IMCBio, part of the ITI 2021-2028 program of the University of Strasbourg, CNRS and Inserm, which was supported by IdEx Unistra (ANR-10-IDEX-0002), and by SFRI-STRAT’US project (ANR 20-SFRI-0012) and EUR IMCBio (ANR-17-EURE-0023) under the framework of the French Investments for the Future Program. DR, KK, and OP are supported by SNSF Sinergia grant CRSII5_183550.

## SUPPLEMENTARY MATERIAL

An online supplement to this article can be found by visiting BJ Online at http://www.biophysj.org.

### List of movies

Movie 1: Actin, myosin and RhoA in a homogeneous giant cell. Time-lapse of actin and Rac densities in the ventral cortex of a giant MDCK cell with no structures. Time is indicated in hh:mm:ss. https://seafile.unistra.fr/smart-link/2d6fb833-35fd-4aea-acc7-88a765ece45c/

Movie 2: Localised state. Time-lapse of the assembly and persistence of a localised state in the cortex of a giant MDCK cell. Time is indicated in hh:mm:ss. https://seafile.unistra.fr/smart-link/9f5dd25a-c431-43ab-bd78-8fb6e6d423d1/

Movie 3: Stress fibres and Rho activity. Time-lapse of actin, active RhoA and myosin densities in the ventral cortex of a giant MDCK cell with fibres. Time is indicated in hh:mm:ss. See snaphots in Suppl. Fig. 7. https://seafile.unistra.fr/smart-link/4eb08d28-490f-4148-a869-c58133af7c88/

Movie 4: Waves actin, myosin, RhoA in giant cells. Time-lapse of actin, active RhoA and myosin densities in the ventral cortex of a giant MDCK cell with waves. Time is indicated in hh:mm:ss. See snaphots in Suppl. Fig. 7. https://seafile.unistra.fr/smart-link/fd9352c1-9b01-43f9-9a3c-5e83895096bf/

Movie 5: Waves actin and Rac localisation. Time-lapse of actin and Rac densities in the ventral cortex of a giant MDCK cell with waves. Time is indicated in hh:mm:ss. See snaphots in Suppl. Fig. 7. https://seafile.unistra.fr/smart-link/1d6645a5-740f-4ea0-8664-c8d2c233df5e/

Movie 6: Cytoplasts and RhoA activation. Time-lapse of an optogenetic RhoA stimulation experiment in a REF52 cytoplast, showing reversible activation of RhoA and reversible contractility of the cytoplast. The blue rectangles indicate position and duration of the stimulation. Time is indicated in hh:mm:ss. https://seafile.unistra.fr/smart-link/1dc30acc-a7e3-4934-95a2-f44f106921fa/

Movie 7: RhoA activation leads to reversible contraction. Time-lapse of an optogenetic RhoA stimulation experiment in a REF52 cell, showing reversible activation of RhoA and reversible contractility. The blue ellipse indicates position and duration of the stimulation. Time is indicated in hh:mm:ss. https://seafile.unistra.fr/smart-link/504963cd-6c54-4a2b-9a9c-49448f4

Movie 8: Spontaneous waves induced in REF52. Time-lapse of actin and active RhoA REF52 cell. Time is indicated in hh:mm:ss. https://seafile.unistra.fr/smart-link/b0906408-0c2b-4a47-b279-acc79afc9031/

Movie 9: RhoA-induced waves in REF52. Time-lapse of a RhoA optogenetic stimulation experiment in a REF52 cell. Optogenetic stimulation is performed in a disk region at times 00:04:20, 00:06:20, 00:08:20, 00:10:20 each time, generating a pulse of RhoA activity followed by a wave of actin and RhoA which expands outward. Time is indicated in hh:mm:ss. https://seafile.unistra.fr/smart-link/0697e133-ace1-4517-bf95-dfc61d125cee/

Movie 10: Waves in the presence of blebbistatin. Time-lapse of a myosin inhibition experiment in a giant MDCK cell. The movie shows a cell untreated (from time 00:00:00 to 01:51:31), and then the same cell treated with blebbistatin 10 μM (from time 01:51:31 to 03:16:11) and 20μM (from time 03:16:11 to 04:32:35). Time is indicated in hh:mm:ss. https://seafile.unistra.fr/smart-link/3254f8f0-8d3b-4137-a3b7-da3d605e1bd1/

Movie 11: Fibres disappear with blebbistatin. Time-lapse of a myosin inhibition experiment in a giant MDCK cell with many fibres. Upon treatment with blebbistatin (starting at time 01:59:58), fibres disappear. Time is indicated in hh:mm:ss. https://seafile.unistra.fr/smart-link/0a8dcf87-2361-45d8-875c-005aa2f3f9fc/

Movie 12: Waves in the presence of Rac1 inhibitor. Time-lapse of a Rac1 inhibition experiment in a giant MDCK cell. The movie shows a cell untreated (from time 00:00:00 to 01:04:07), and then the same cell treated with Rac1 inhibitor II 100 μM (from time 01:04:07 to 03:12:22). Time is indicated in hh:mm:ss. https://seafile.unistra.fr/smart-link/aaad6c15-3e44-47c1-b882-d85c8f7c69cd/

Movie 13: FRAP of Rac1. Time-lapse of a FRAP experiment of Rac1 in a giant MDCK cell, in the front of a wave. The wave is visible with both actin and Rac1 channels, moving from the bottom to the top. Rac1 is photobleached twice (at time 00:00:53 and at time 00:01:24) in the wave front, and recovers within seconds. Time is indicated in hh:mm:ss. https://seafile.unistra.fr/smart-link/2d226136-9e25-47f3-8859-7cc9a6b4eab0/

Movie 14: Actin waves generate mechanical forces. Time-lapse of fiducial markers in a giant MDCK cell while waves are passing. Fiducial markers (visible in actin fluorescence and transmitted light) move and their trajectory correlates with the direction of waves, indicating that waves generate mechanical forces in the cortex. Time is indicated in hh:mm:ss. https://seafile.unistra.fr/smart-link/c76d2064-a350-4cb3-869b-9e735a0ce5a7/

Movie 15: Waves push fiducial markers. Time-lapse on actin waves in giant cells pushing fiducial markers. Fiducial markers tracks are displayed in blue. Time is indicated in hh:mm:ss. https://seafile.unistra.fr/smart-link/2690daba-48b9-46ee-b0a4-3c

Movie 16: Wave transforms into lamellipodia when reaching the cell edge. Time-lapse of an actin wave reaching the edge of a giant MDCK cell. The wave transforms into a lamellipodia. Time is indicated in hh:mm:ss. https://seafile.unistra.fr/smart-link/15f51b81-3ee3-41b8-9942-f5c2585d9438/

Movie 17: Numerical simulation of waves. https://seafile.unistra.fr/smart-link/ae920b49-fffc-417e-943d-bdf0dda2d22

Movie 18: Numerical simulation of waves: interference. Simulation of two waves meeting and annihilating. Simulation parameters are: … https://seafile.unistra.fr/smart-link/d21c93c7-4144-43a7-b900-94d1b023d64e/

Movie 19: Numerical simulation waves: Rac inhibition. Numerical simulation of a Rac inhibition / RhoA activation experiment. Rac is inhibited in a disk region, and a wave propagates to this region. When entering the region, the waves annihilates. https://seafile.unistra.fr/smart-link/a0a5a449-7291-4012-a365-659b60b5e569/

Movie 20: Interference between actin waves. Time-lapse of the interaction of two actin waves which meet front to front in a giant MDCK cell. The wave fronts annihilate. Time is indicated in hh:mm:ss. https://seafile.unistra.fr/smart-link/9de20196-1565-4250-8fb6-94aeb9726aa7/

Movie 21: Optogenetic activation of Rac1. Time-lapse of a Rac1 optogenetic stimulation experiment in the ventral cortex of a giant MDCK. The blue disk indicates position and duration of the optogenetic stimulation. Upon release of stimulation (blue disk disappears), the cortex suddenly collapses and relaxes elastically, indicating that a tension was generated in the cortex by the stimulation. Time is indicated in hh:mm:ss. https://seafile.unistra.fr/smart-link/21c5ffde-3b62-4ef6-afeb-f0943b2e624b/

Movie 22: Time-lapse of a Rac1 optogenetic stimulation experiment in the ventral cortex of a giant MDCK. The blue disk indicates position and duration of the optogenetic stimulation. Upon release of stimulation (blue disk disappears), a wave of actin propagates radially outward from the disk, to the lower right direction. Time is indicated in hh:mm:ss. https://seafile.unistra.fr/smart-link/3e767004-da81-4a1e-9ba1-b6a0d87aa41c/

Movie 23: Optogenetic activation of RhoA can be ignored by actin waves. Time-lapse of a RhoA optogenetic stimulation experiment in a giant MDCK cell at the front of a wave. The blue rectangle indicates position and duration of the optogenetic stimulation, at the wave front. The wave continues to propagate unaltered. Time is indicated in hh:mm:ss. https://seafile.unistra.fr/smart-link/419598ed-9a6b-43a8-aa6f-646c067cb10e/

Movie 24: Optogenetic activation of RhoA can slow actin waves. Time-lapse of a RhoA optogenetic stimulation experiment in a giant MDCK cell at the front of a wave. The blue rectangle indicates position and duration of the optogenetic stimulation, at the wave front. The portion of wave intercepted by the stimulation slows down, curving the wave front. Time is indicated in hh:mm:ss. https://seafile.unistra.fr/smart-link/fc0d65fa-09bd-4d59-9337-a0e67ea29375/

Movie 25: Optogenetic activation of RhoA can break actin waves. Time-lapse of a RhoA optogenetic stimulation experiment in a giant MDCK cell at the front of a wave. The blue ellipse indicates the position and duration of the optogenetic stimulation, at the wave front. The portion of wave intercepted by the stimulation breaks, yielding two wave fronts which propagate in opposite directions. Time is indicated in hh:mm:ss. https://seafile.unistra.fr/smart-link/03f3c606-d9ce-4cd1-aa09-e5ce41416b77/

### List of supplementary figures

SF1: Diameter of giant cells over time during cytochalasin D culture. Growth curve of giant MDCK cell, during culture with cytochalasin D incubation (see Methods). Cells were measured before (blue) and after (orange) 1h washout of the cytochalasin D. https://seafile.unistra.fr/smart-link/d5f84e3f-d1b2-44e8-ace4-62dd08ba9bab/

SF2: Patterns are similar in transfected and in non-transfected cells. Illustration of fibres (top) and waves (bottom) in giant wild-type MDCK cells. Actin, myosin and nuclei were stained with anti-bodies or DAPI (see Methods). Fibres and waves are similar to that in transfected cells. https://seafile.unistra.fr/smart-link/2e0d318d-ccbe-45cd-9ac6-1184bef27c4a/

SF3: Snapshots of Movies 3, 4 and Movie 5, showing waves and fibres, respectively. https://seafile.unistra.fr/smart-link/b94863a9-5586-4de6-9f7e-08372ec5429d/

SF4: 3D representation of a giant MDCK cell, with a wave. Left: XY, YZ, XZ projections of actin in a giant MDCK cell, the aspect ratio of the cell is very large and the cell cortex is very flat (except where the nucleus are). Right: zoom on the wave (yellow box on left panel) with corresponding projections. The wave moves from the top to the bottom, the ventral membrane is not flat at the front of the wave as shown on the YZ projection. https://seafile.unistra.fr/smart-link/73ebd6c7-7232-4c34-b152-ee1f8844792e/

SF5: Fibres in a REF52 and in a cytoplast. Top: Actin and RhoA activity in a REF52 cell with many stress fibres. Bottom: Actin and DAPI (nucleus) in a cytoplast of a REF52. Both systems show many fibres, indicating genericity of this structure. https://seafile.unistra.fr/smart-link/9e004d08-56f7-4e88-a22e-a08232f5ae42/

SF6 : Spontaneous Actin/RhoA waves in PDGF-Stimulated REF52 fibroblasts. (A) Overview of a REF52 cell imaged 48 hours after stimulation with 50 ng/ml PDGF. The yellow box denotes the region shown in (B), and the red line indicates where the kymograph was generated in (C). Scale bar, 25 μm.(B) Time-lapse montage (0–8 minutes) of a single propagating wave. Top row: Actin (Lifeact::mNeonGreen). Middle row: Active RhoA (2xrGBD::dTomato). Bottom row: Merged channels (Actin in green, Active RhoA in magenta). Scale bar, 5 μm. (C) Kymograph along the red line in (A). Top panel: Actin (Lifeact::mNeonGreen). Middle panel: Active RhoA (2xrGBD::dTomato). Bottom panel: Merged channels. The yellow line in the composite panel marks the region used for the intensity profile in (D). Scale bar, 10 μm (vertical) and 20 minutes (horizontal). (D) Normalized intensity profiles of Actin (green) and Active RhoA (purple) measured along the yellow line in (C), plotted over time. To correct for photobleaching, all images underwent histogram matching. Rigid-body registration was then performed using pystackreg. For the intensity measurements in the plot, each set of values was normalized by subtracting the mean and subsequently scaled by dividing by the standard deviation. See also Movie 8. https://seafile.unistra.fr/smart-link/c1767565-4c52-4f4f-ac31-cbd3550eb061/

SF7: Waves in REF52 stimulated by optogenetic activation of RhoA (Movie 9). (A) Left: Time series of optogenetic stimulation of RhoA in a REF52 cell. The blue disk indicates the position and time of the optogenetic stimulation which is maintained for one second. After activation, a wave of actin and Rac propagates outward from the disk region. Right: overview of the same cell; the yellow box indicates the region that is enlarged on the left panel. (B) Overview of a REF52 cell showing Actin (Lifeact::mNeonGreen) and active RhoA (2xrGBD::dTomato), box denotes the region shown in (C).(C) Timelapse imaging shows Actin and RhoA recruitment after a weak optogenetic stimulation with optoLARG in circular region. (D) Same timeseries as in (B), but with the mean of the first 20 acquired frames subtracted to highlight the change in biosensors.(E) Averaged intensity profile of 72 stimulations in n=36 cells at stimulation location shows dynamics of actin recruitment and RhoA activation after stimulation. (F) Radially averaged kymographs of same experiment as in (E) shows propagation of signal in space. In these experiments, cells are segmented using an interactive pixel classifier [https://doi.org/10.1101/2024.09.12.610926] and two stimulation locations are automatically chosen for stimulation along the major axis of the cell using a custom python script. https://seafile.unistra.fr/smart-link/60915ca2-33ee-49a6-8708-b6ee4635f270/

SF8: Wave speed with blebbistatin and Rac1 inhibitor II. Measurements of wave speed in various experimental conditions. https://seafile.unistra.fr/smart-link/9d98d03d-b14a-4279-8e3e-1c4d4291eb58/

