## Supplementary figures and images for "Interplay between Rac1/RhoA and actin waves in giant epithelial cells : experiment and theory"

### SF1

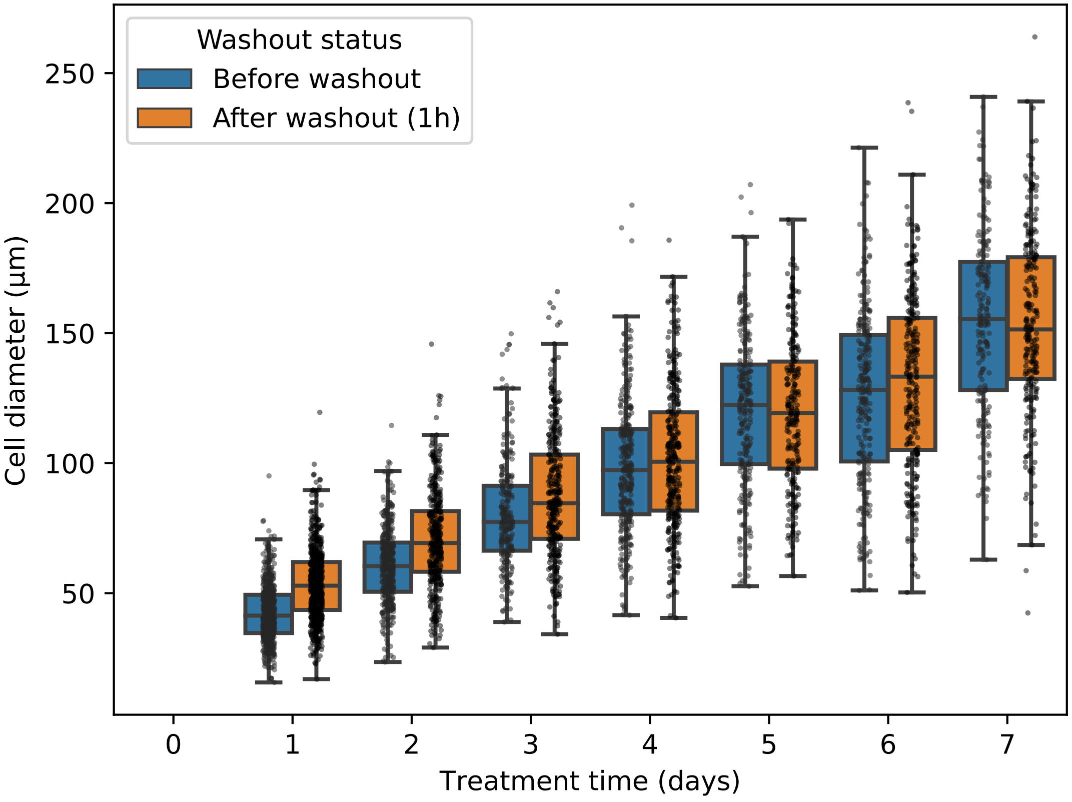

### SF2

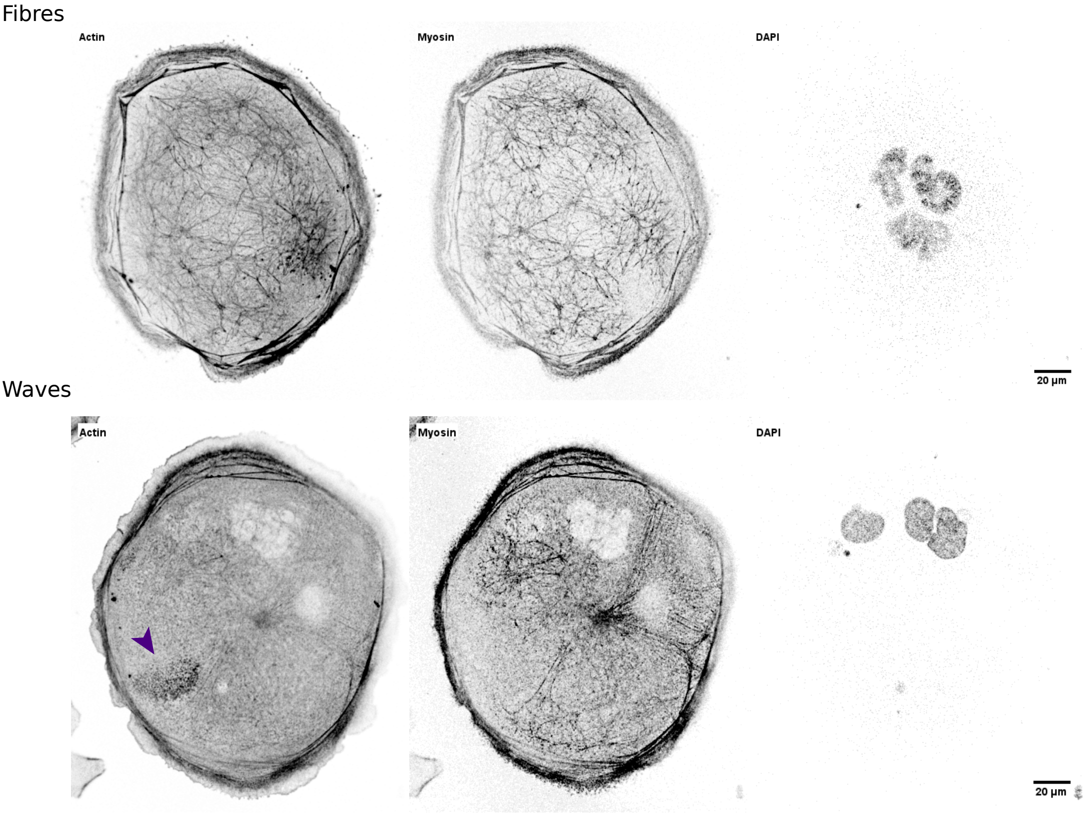

### SF3

#### Waves

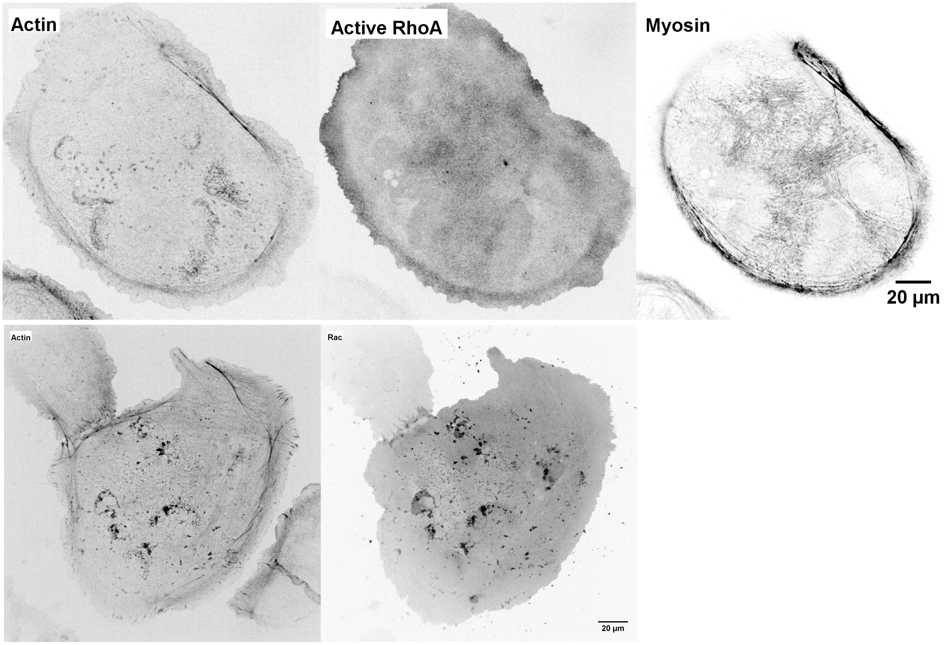


#### Fibres

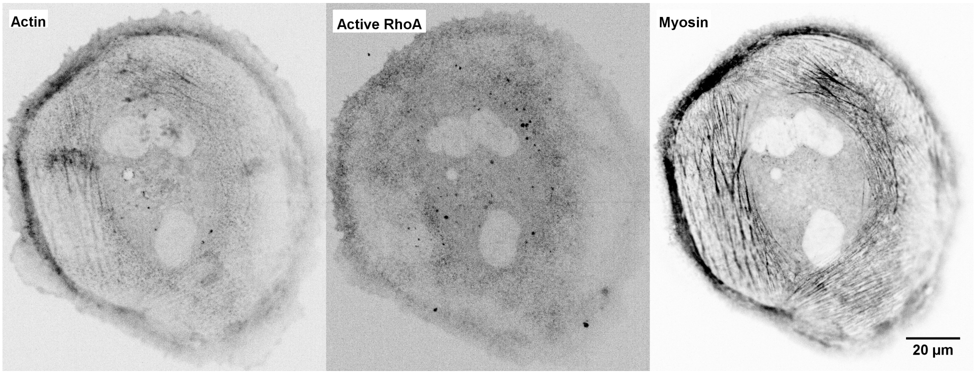

### SF4

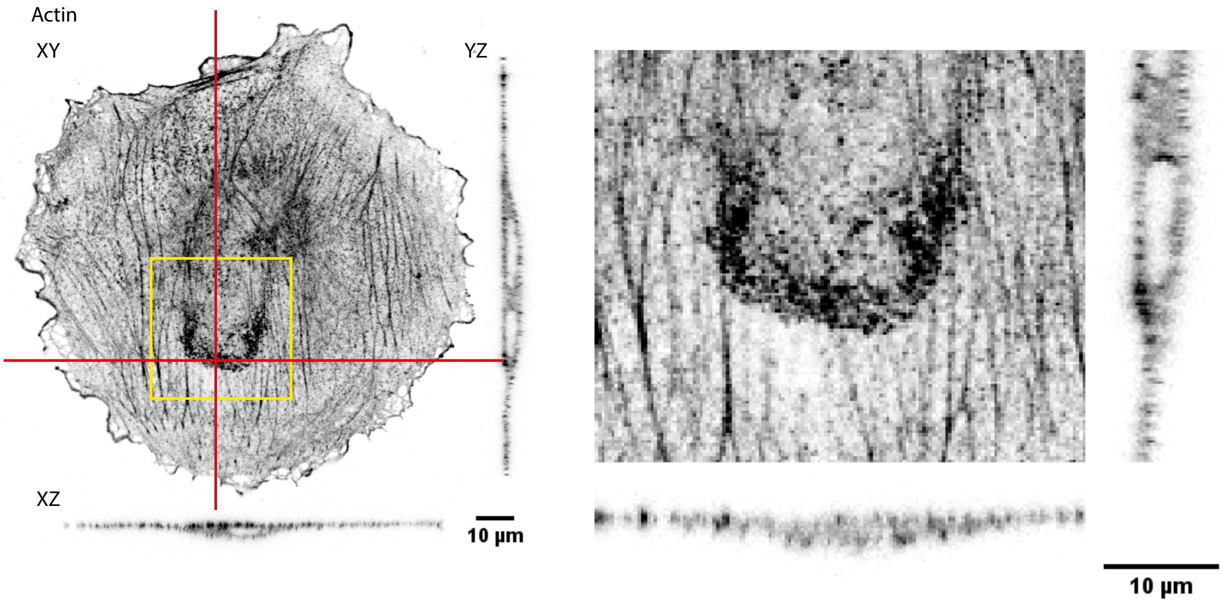

### SF5

REF52

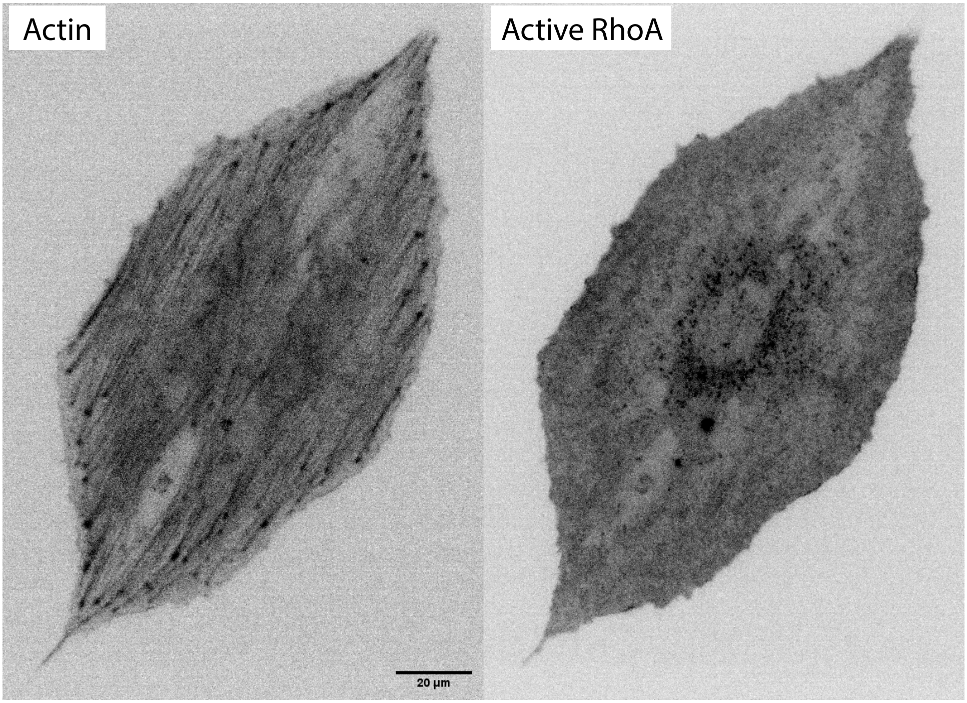


Actin

DAPI

Cytoplast of REF52

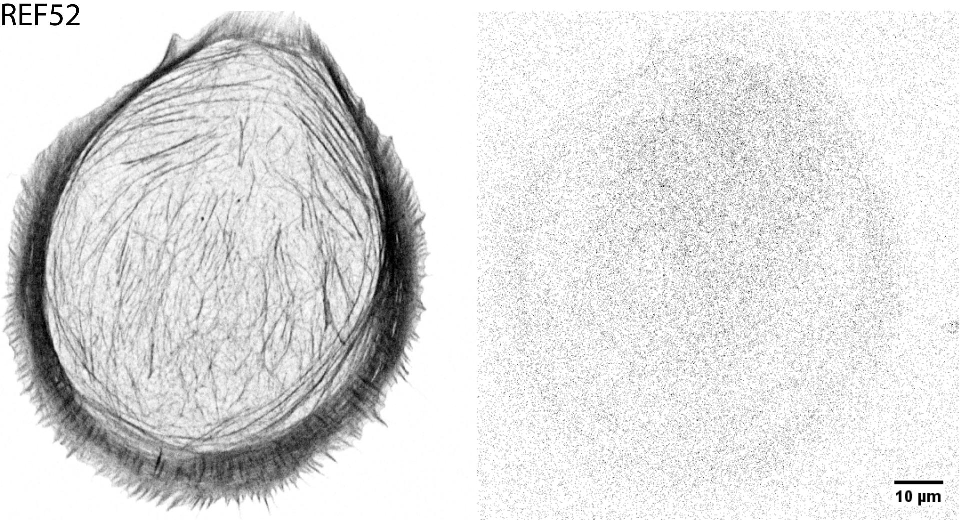

### SF7

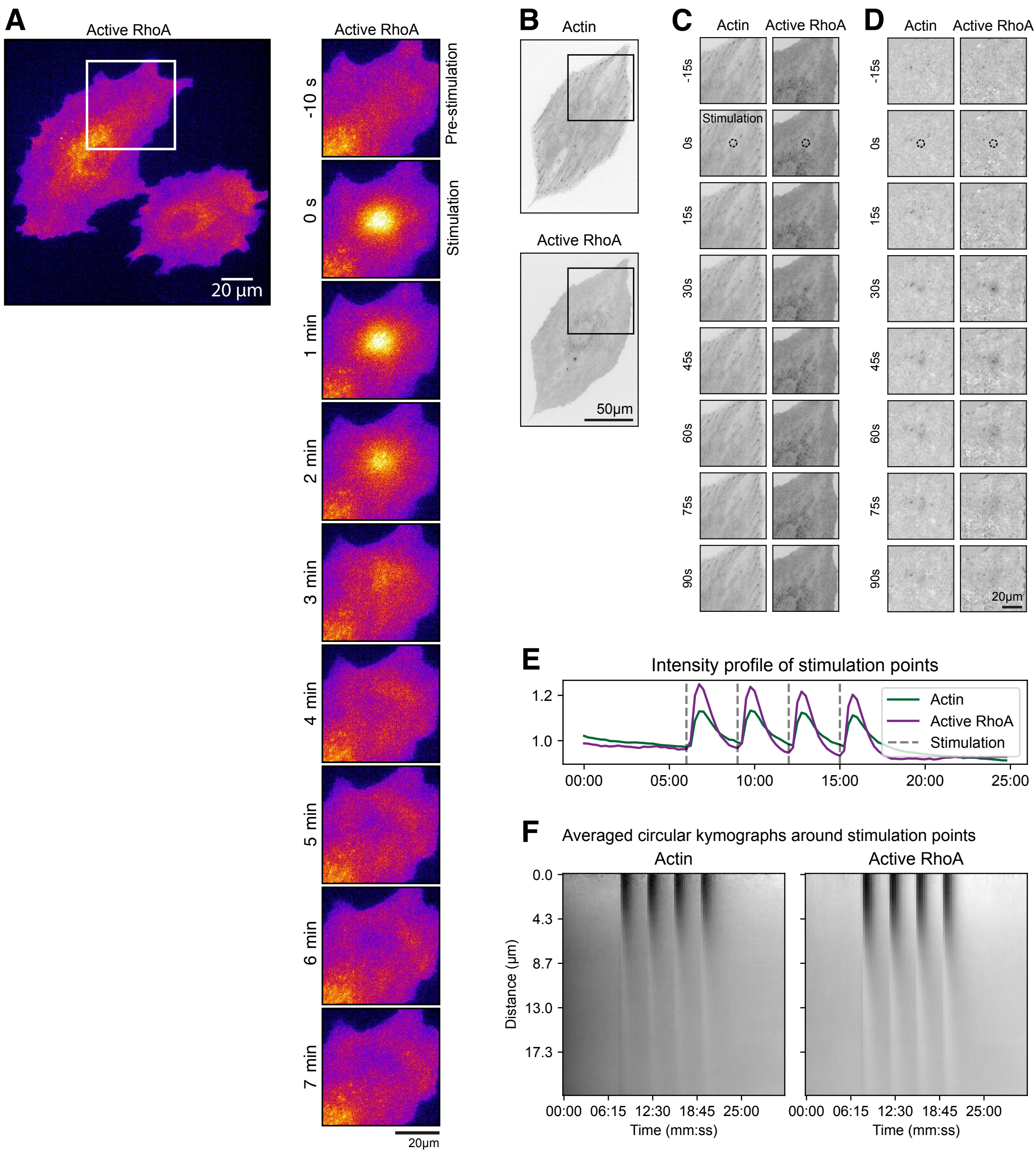

### SF8

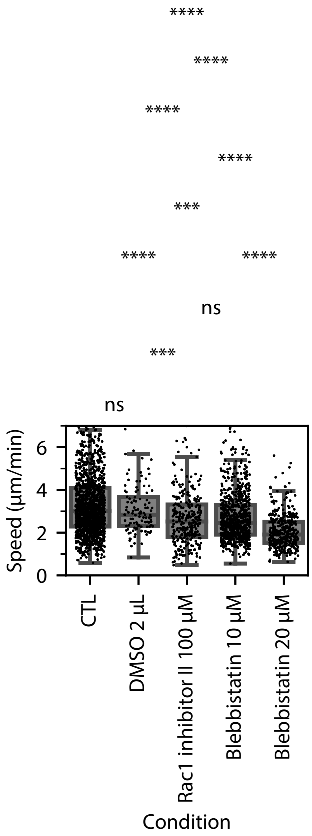
