## Supplementary material for "Interplay between Rac1/RhoA and actin waves in giant epithelial cells : experiment and theory": SF6

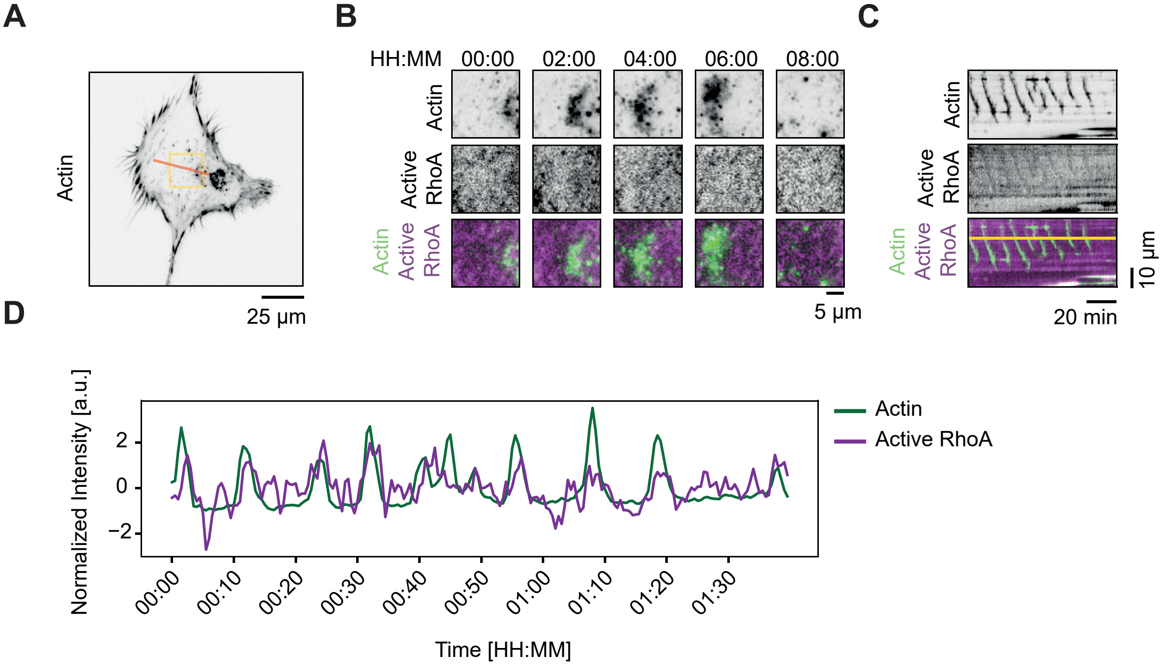


Spontaneous Actin/RhoA Waves in PDGF-Stimulated REF52 Fibroblasts.
(A) Overview of a REF52 cell imaged ~48 hours after stimulation with 50 ng/ml PDGF.
The yellow box denotes the region shown in (B), and the red line indicates where the kymograph in (C) was generated. Scale bar, 25 µm.
(B) Time-lapse montage (0–8 minutes) of a single propagating wave. Top row: Actin (Lifeact::mNeonGreen).
Middle row: Active RhoA (2xrGBD::dTomato). Bottom row: Merged channels (Actin in green, Active RhoA in magenta). Scale bar, 5 µm.
(C) Kymograph along the red line in (A). Top panel: Actin (Lifeact::mNeonGreen). Middle panel: Active RhoA (2xrGBD::dTomato).
Bottom panel: Merged channels. The yellow line in the composite panel marks the region used for the intensity profile in (D).
Scale bar, 10 µm (vertical) and 20 minutes (horizontal).
(D) Normalized intensity profiles of Actin (green) and Active RhoA (purple) measured along the yellow line in (C), plotted over time.
